# Mechanical stress orients stomata division to form tissue scale alignments

**DOI:** 10.1101/2024.12.02.626480

**Authors:** Leo Serra, Euan T. Smithers, Lucy Bentall, Martin O. Lenz, Sarah Robinson

## Abstract

The last stomatal division aligns with the leaf’s main axis in many species [1]. Understanding how cellular events such as these are coordinated across organ scales remains a challenge in developmental biology. In Arabidopsis, polarised proteins guide the asymmetric divisions in the early stomatal lineage. These proteins show organ scale alignment and may be sensitive to mechanical stress [2]. In contrast, what determines the orientation and alignment of the critical final division is unknown [3]. Here we use an artificial system where every cell adopts the fate of a stomata pore [4] making it easy to visualise their alignment. Combining this system with simultaneous time-lapse imaging on both sides of the cotyledon we are able to compare the stomatal orientation relative to the organ axis, the cell major axis, and the principal directions of growth. Using finite element modelling on a realistic template enabled us to identify differential growth-derived stress patterns as a factor coordinating stomata division at the organ scale. Mechanical perturbation confirmed the influence of tensile stress on stomata division orientation. Through this study, we have identified a mechanism that can explain this nearly century-old observation.

## Results

### Orientation of stomata changes through development and differs between the abaxial and adaxial sides of the cotyledon

To characterise the pattern of stomata orientation during cotyledon development we manually labeled more than 10000 stomata on 72 cotyledons from 1 to 5 days after germination (DAG), their orientation was then mapped relative to the cotyledons proximo-distal axis (PD axis) (Figure 1). On the abaxial side, most of the stomata are aligned with the PD axis at 1 DAG (Figure 1A), and over 40% were within 10 degrees of the PD axis (Figure 1B). At 5 DAG the stomata are still preferentially aligned with the PD axis but to a lesser extent with only 20% of stomata diverging from the PD axis by less than 10 degrees (Figure 1C). This indicates that the newly formed stomata are not aligned with the PD axis anymore. On the adaxial side, the first stomata formed at 1 DAG are aligned with the PD axis, but as soon as 2 DAG the new stomata diverge from this axis (Figure 1D and E). These differences in orientations between early and late developmental stages as well as between abaxial and adaxial sides means that the factor coordinating stomata division across the organ must be changing between these contexts. Since the difference of pattern was evident between the two sides of the cotyledons at 2 DAG and young samples are more convenient to work with, we decided to focus on the abaxial adaxial differences occurring before 2 DAG for the rest of our study.

**Figure 1:**
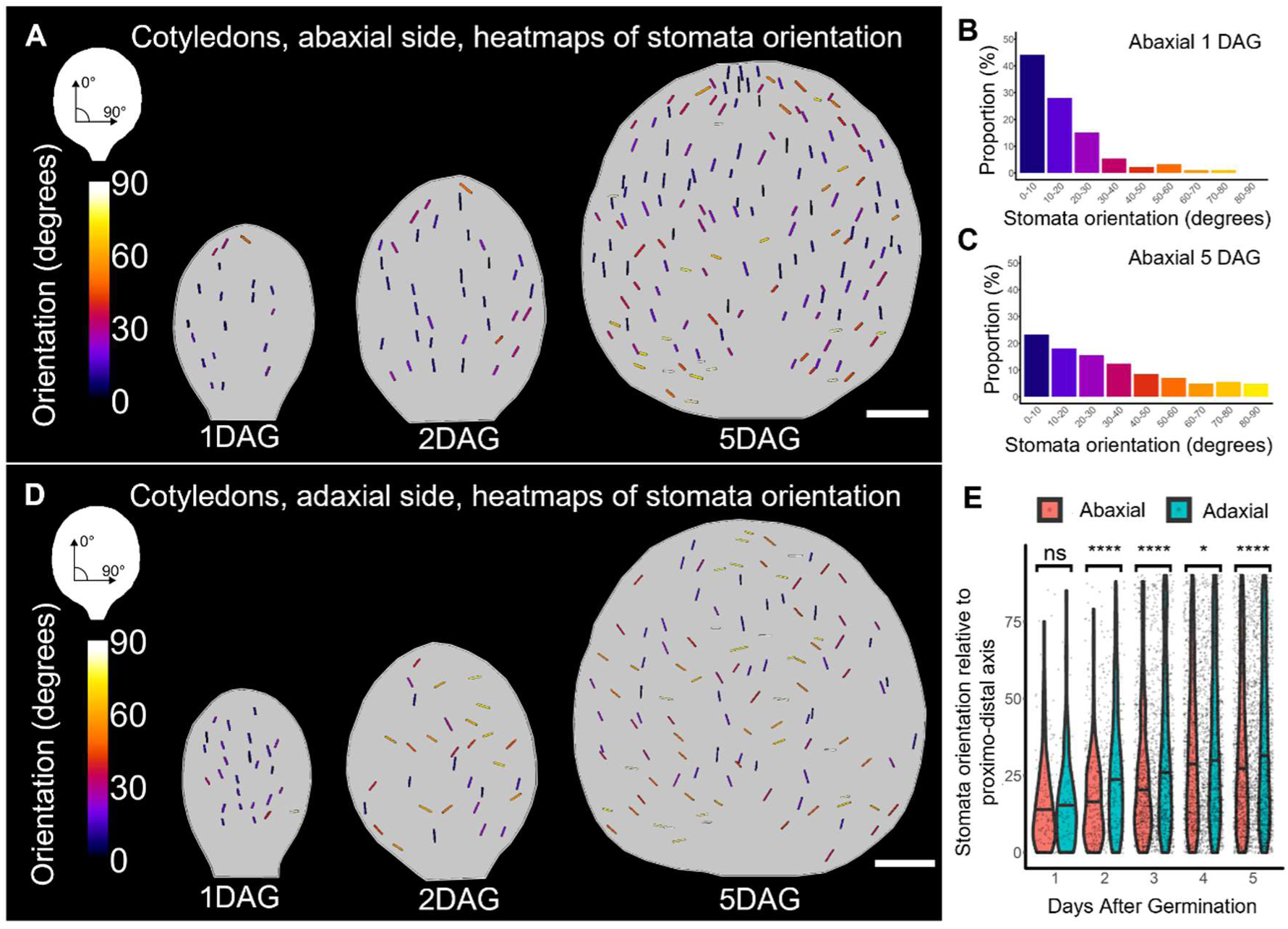
Orientation of stomata changes through development and differs between the two sides of the cotyledon. (A) Map of stomata orientation relative to proximo-distal axis on the abaxial sides of cotyledons (each line represents a stomata, scale bar:200μm). (B) Proportion of different stomata orientations relative to proximo-distal axis on the abaxial side of cotyledons at 1 DAG (93 stomata from 5 cotyledons). (C) Proportion of different stomata orientations relative to proximo-distal axis on the abaxial side of cotyledons at 5 DAG (1154 stomata from 7 cotyledons). (D) Map of stomata orientation relative to proximo distal axis on the adaxial sides of cotyledons (each line represents a stomata, scale bar:200μm). (E) Distributions of stomata orientations relative to proximo-distal axis on both sides of cotyledons during early developmental stages (abaxial side 5899 stomata from 33 cotyledons; adaxial side 4402 stomata from 39 cotyledons).

### Organ scale coordination of stomata division orientation is independent of cell growth direction and cell geometry

Cell growth and cell geometry have long been identified as the main factors directing cell division orientation [5–8], we thus wanted to investigate their contributions to the orientation of stomata division. To do so we took advantage of the MUTE estradiol inducible line [4] which can trigger stomata differentiating division in every shoot epidermal cell. We crossed it with a plasma membrane marker [9] (iMUTExPM-YFP hereafter), performed time lapse imaging on the abaxial and adaxial side of cotyledons, and analysed the division orientation (Figure 2). When MUTE was induced at germination (0DAG), it resulted in the formation of divisions that showed good alignment with the PD axis on both sides of the cotyledons (Figure 2A). However, when the divisions were induced at 1 DAG, the alignment with the PD axis was still clear on the abaxial side but not on the adaxial side (Figure 2B). To evaluate if the induced divisions were directed by cell geometry, cell growth, or an organ-scale derived factor, we analysed these time-lapses images with MorphoGraphX [10]. Cell segmentation and lineage tracking allowed us to identify the clones (group of sister cells deriving from a single mother cell at t0) resulting from a single division event, we then extracted these clones’ principal growth, and major axis, as well as cotyledon PD axis, we then compared the division orientation with these axes (Figure 2C). When the divisions were induced at 0DAG they were very close to the cell major axis and organ PD axis on both sides of the cotyledon with 50% of the analysed division diverging with these axes by less than 15 degrees, at the same time, the divergence with clone growth axis was more than 25 degrees on the abaxial side and more than 50 degrees on the adaxial side for half of the divisions (Figure 2D). For divisions induced at 1DAG, the divergence with the clone major axis was still low on both sides of the cotyledon (less than 25 degrees for 50% of the divisions), and the divergence with the growth axis was even higher than at 0DAG (more than 50 degrees of divergence for most of the divisions). The relationship with the PD axis was more contrasted: while on the abaxial side, the divisions were still aligned with the PD axis (half of the divisions within 20 degrees of the PD axis), on the adaxial side they were less aligned with this axis (50% of the divisions within 25 degrees of the PD axis) (Figure 2E). These first analyses allowed us to exclude growth direction as a factor guiding stomata division orientation, however, it was not possible to discriminate between clone geometry and PD axis.

**Figure 2:**
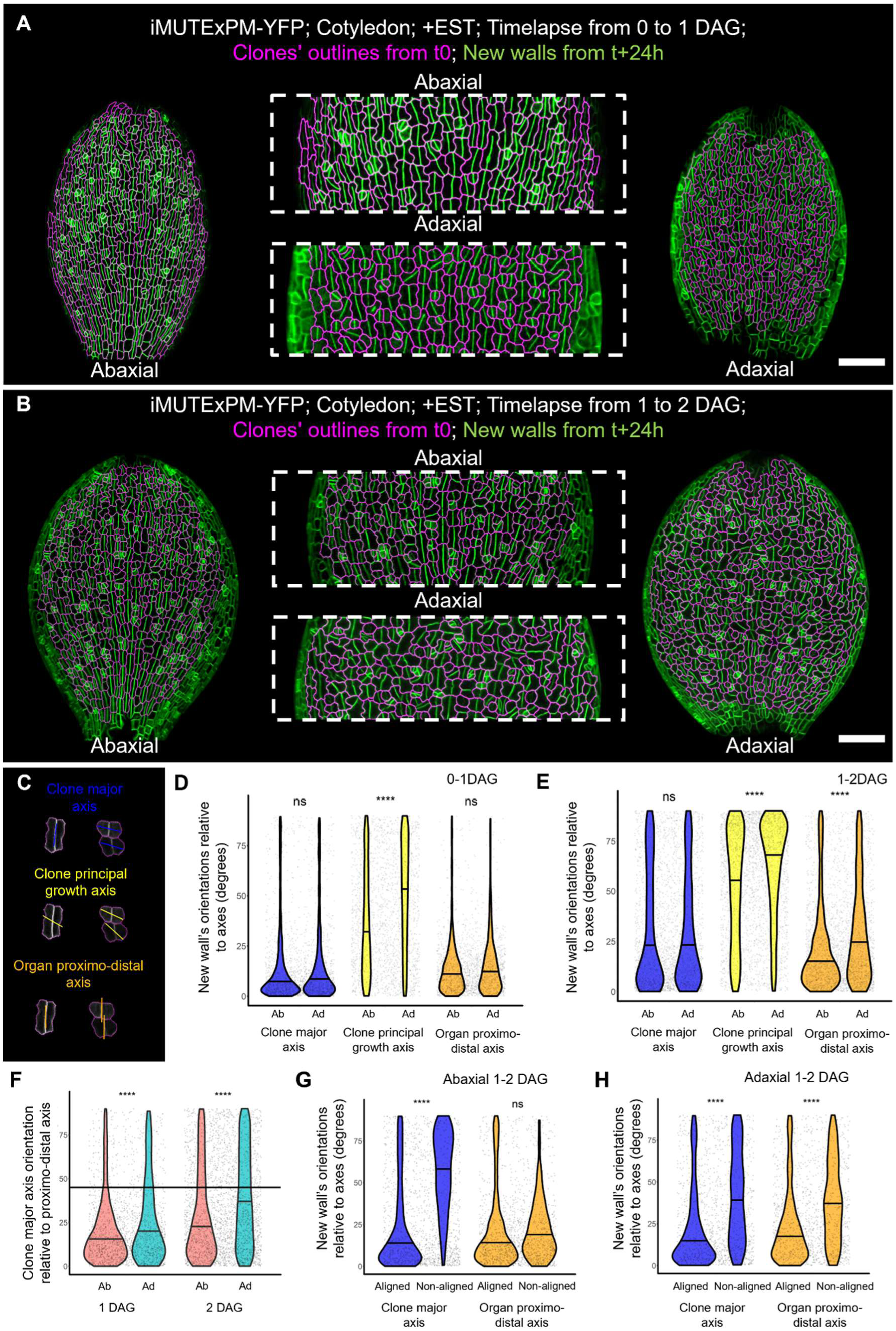
Stomata division orientations along the proximodistal axis differ between the two sides of the cotyledons and are independent of cell growth and geometry. (A) Stomata differentiating divisions induced between 0 and 1 DAG on the abaxial and adaxial side of iMUTExPM-YFP cotyledons (Clones from t0 are outlined in magenta, the surface at t+24h highlighting the new walls is in green. Scale bar:100μm). (B) Stomata differentiating divisions induced between 1 and 2 DAG on the abaxial and adaxial side of iMUTExPM-YFP cotyledons (Clones from t0 are outlined in magenta, the surface at t+24h highlighting the new walls is in green. Scale bar:100μm). (C) Visualisation and quantification of induced stomata-differentiating divisions: for each clones, the orientation of the new wall is compared to the orientation of the clone major axis (blue lines), the clone principal growth axis (yellow lines), or the organ proximo-distal axis (orange lines). (D) Distribution of the orientation of new walls relative to the clone major axis, clone principal growth axis, or organ proximo-distal axis for divisions induced between 0 and 1 DAG (data for abaxial time-lapses comes from 3 independent experiments and a total of 1434 divisions analysed, data for adaxial time-lapses come from 3 independent experiments and a total of 547 divisions analysed). (E) Distribution of new walls orientations relative to clone major axis, clone principal growth axis, or organ proximo-distal axis for divisions induced between 1 and 2 DAG (data for abaxial time-lapses comes from 4 independent experiments and a total of 1575 divisions analysed, data for adaxial time-lapses come from 4 independent experiments and a total of 1356 divisions analysed). (F) Distribution of the orientations of clones’ major axis relative to organ proximo distal axis (horizontal line marks the 45 degrees threshold used to define clones aligned or not aligned with the organ proximo distal axis). (G) Distribution of new walls orientations relative to clone major axis or organ proximo-distal axis for clones aligned/not aligned with organ proximo distal axis on timelapses on the abaxial side between 1 and 2 DAG (1117 aligned divisions and 398 non-aligned divisions). (H) Distribution of new walls orientations relative to clone major axis or organ proximo-distal axis for clones aligned/not aligned with organ proximo distal axis on timelapses on the adaxial side between 1 and 2 DAG (797 aligned divisions and 559 non-aligned divisions).

To discriminate between cell geometry and organ scale factors, we quantified the orientation of cell major axes relative to PD axes on both sides of cotyledons at 0 and 1 DAG (Figure 2F), and defined non-aligned and aligned clones based on a 45 degrees threshold of divergence between organ axis and clone major axis (horizontal line in Figure 2F). We then compared the orientation of division induced at 1 DAG relative to cell geometry or organ axis for aligned versus non-aligned clones (Figure 2G and 2H). On the abaxial side, while the division alignment with the organ axis remained for cells not aligned with the PD axis, the alignment with the clone major axis was lost (Figure 2G). On the adaxial side, the alignment with both clone major axis and organ axis was disrupted for the clones not aligned with the PD axis (Figure 2H). Altogether these results suggest that the orientation of stomata divisions is independent of cell geometry and growth. These results also revealed that while the division where aligned with the PD axis on the abaxial side of the cotyledon, they were more disorganised on the other side. To further identify the causative factor coordinating these divisions we decided to first characterise the differences between the abaxial and adaxial sides of the cotyledons in more detail.

### Differential growth between the two sides of the cotyledon generates orthogonal mechanical stress patterns

To get a better picture of changes taking place between both sides of the cotyledon during early development, light sheet imaging was used to get a full 3D view of cotyledons at 1 and 2 DAG (Figure 3A). These images and the virtual transverses sections (Figure 3B and C) revealed drastic changes of curvature between 1 and 2 DAG: at 1DAG the adaxial epidermis is flat and the abaxial epidermis is convex, at 2 DAG the adaxial epidermis become convex and the abaxial epidermis become concave. Such changes might result from differential growth between the two sides. To evaluate this, we performed dual-view timelapse imaging to map growth on both sides of the same cotyledons (Figure 3D). The analysis confirmed that the two sides underwent differential growth with growth ratio (clone area at t=24h/mother cell area at t0h) from 1 to 2DAG being around 3 for the adaxial side and around 2 for the abaxial side (Figure 3D and E). To determine if differential growth rates can generate different stress patterns on the two sides of the cotyledon we used a finite element modeling approach. Geometric data extracted from a 3D segmented cotyledon at 1DAG (Figure 3F) was used to generate a realistic cotyledon cross-section mesh containing adaxial, and abaxial epidermis, mesophyll and palisade cells (Figure 3G). The total curvature and aspect ratio of the mesh as well as the relative cell widths and heights were maintained. The model was inflated and then allowed to grow using varying growth rates throughout the tissue (Figure 3H). After differential growth, the cotyledon mesh resembled an actual cross-section of a cotyledon at 2 DAG convex on the top and concave on the bottom (Figure 3 C and I). The main direction of the resulting stress was perpendicular to the PD axis of the cotyledon on the top (adaxial side), and parallel to it on the bottom (abaxial side). To experimentally confirm the model prediction we imaged cotyledons of *quasimodo2-2* (Figure 3J), a cell adhesion defect mutant forming cracks between epidermal cells, the localisation of those cracks having been previously used to infer the main direction of tensile stress in plants epidermis [11]. At 2DAG the abaxial side of *quasimodo2-2* had a lot of transverse cracks indicating a tensile stress parallel to the PD axis of the cotyledon, the adaxial side had very few cracks located on the transverse wall indicating a potential tensile stress perpendicular to the PD axis. Taken together these experiments and simulations indicate that a differential growth between the two sides of the cotyledons is associated with orthogonal patterns of tensile stress.

**Figure 3:**
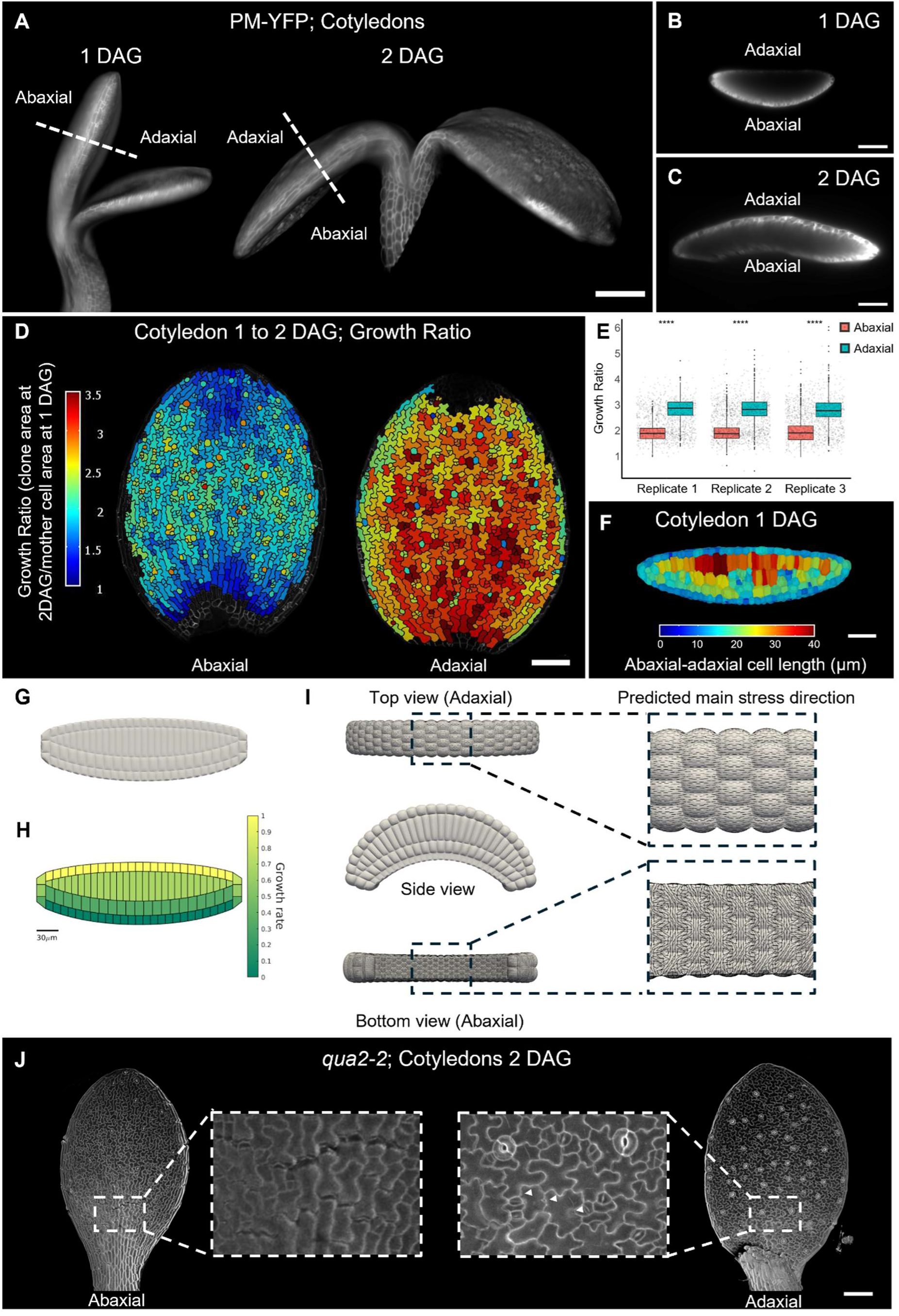
Differential growth between the two sides of the cotyledon generates orthogonal mechanical stress patterns. (A) Light-sheet images of seedlings at 1 and 2 DAG during cotyledons opening (dashed lines indicate the emplacement of optical transverse sections shown in B and C. scale bar: 200μm). (B) Optical transverse section of a cotyledon at 1 DAG (scale bar:100μm). (C) Optical transverse section of a cotyledon at 2 DAG (scale bar:100μm). (D) Heatmaps showing the distribution of growth ratio on the two sides of the same cotyledon time-lapsed between 1 and 2 DAG (scale bar:100μm). (E) Distribution of growth ratio for both sides of the cotyledon presented in D (replicate 1), and two additional replicates. (F) Heatmap showing the distribution of abaxial adaxial cell length in a transverse section of cotyledon at 1 DAG (based on 3D segmentation of a whole cotyledon, scale bar:50μm). (G) A realistic 3D mesh of a section of a cotyledon at 1 DAG based on the geometric data extracted from the sample presented in F. (H) Distribution of the set growth rate used for the simulation in (I). (I) Prediction of the orientation of differential growth-based stress on the two sides of a cotyledon mesh. (J) Different localisation of cell adhesion defect on the 2 sides of cotyledons in a qua2-2 mutant at 2 DAG (arrowheads point to small cracks on the adaxial epidermis).

### Changes in the tensile stress pattern shift the orientation of stomata divisions

To determine if mechanical stress can orient stomata we applied a range of mechanical perturbations. Cutting the tissue or making holes has been used previously to change the direction of maximal tensile stress by releasing tissue tension [12–14]. Here we show that MUTE-induced divisions occur in a different orientation following the cell ablations or tissue cutting in a way that is consistent with them aligning with the direction of maximal tensile stress (Figure S2). To prevent the unavoidable wounding response associated with these approaches we have developed a new protocol to change the mechanical stress pattern in intact cotyledons. Using tape (tough tag©) we have been able to fold a 2 DAG cotyledon on itself, resulting in a drastic change of cotyledon curvature (Figure 4A, B, and C) which is likely to generate tension parallel to the PD axis. To determine the stress pattern in the cotyledon after folding, we modified the model template generated above (Figure 3) to make it longer along the PD axis, we then applied a 3-point bending which was able to produce a folding comparable to the experiment (Figure 4D). The predicted stress pattern in the epidermis differs between the two simulations with the non-folded one having tension oriented perpendicular to the PD axis and the folded one having tension parallel to the PD axis (Figure 4E). Since microtubules have been reported to reorient upon mechanical stress [12–15], we imaged non-folded and folded adaxial sides of cotyledons of 35S::MBD-GFP microtubules reporter to validate experimentally the effect of the folding on the mechanical stress pattern (Figure 4F). 24 hours after folding, cotyledons exhibited an overall orientation of microtubules parallel to the PD axis, at the same time, the non-folded ones had an overall microtubules orientation perpendicular to the PD axis (Figure 4F and H), confirming that the folding was changing the tensile stress pattern. To test if tensile stress was directing stomata division we induced MUTE division in folded and non-folded cotyledons and quantified the orientation of division (Figure 4G and I). When divisions were induced in a non-folded cotyledon, they were mostly oriented transversely to the PD axis, but when they were induced in folded cotyledons, their orientations were mostly parallel to the axis of applied tension (the PD axis). Altogether these experiments demonstrate that tensile stress can coordinate the orientation of stomata division at the organ scale.

**Figure 4:**
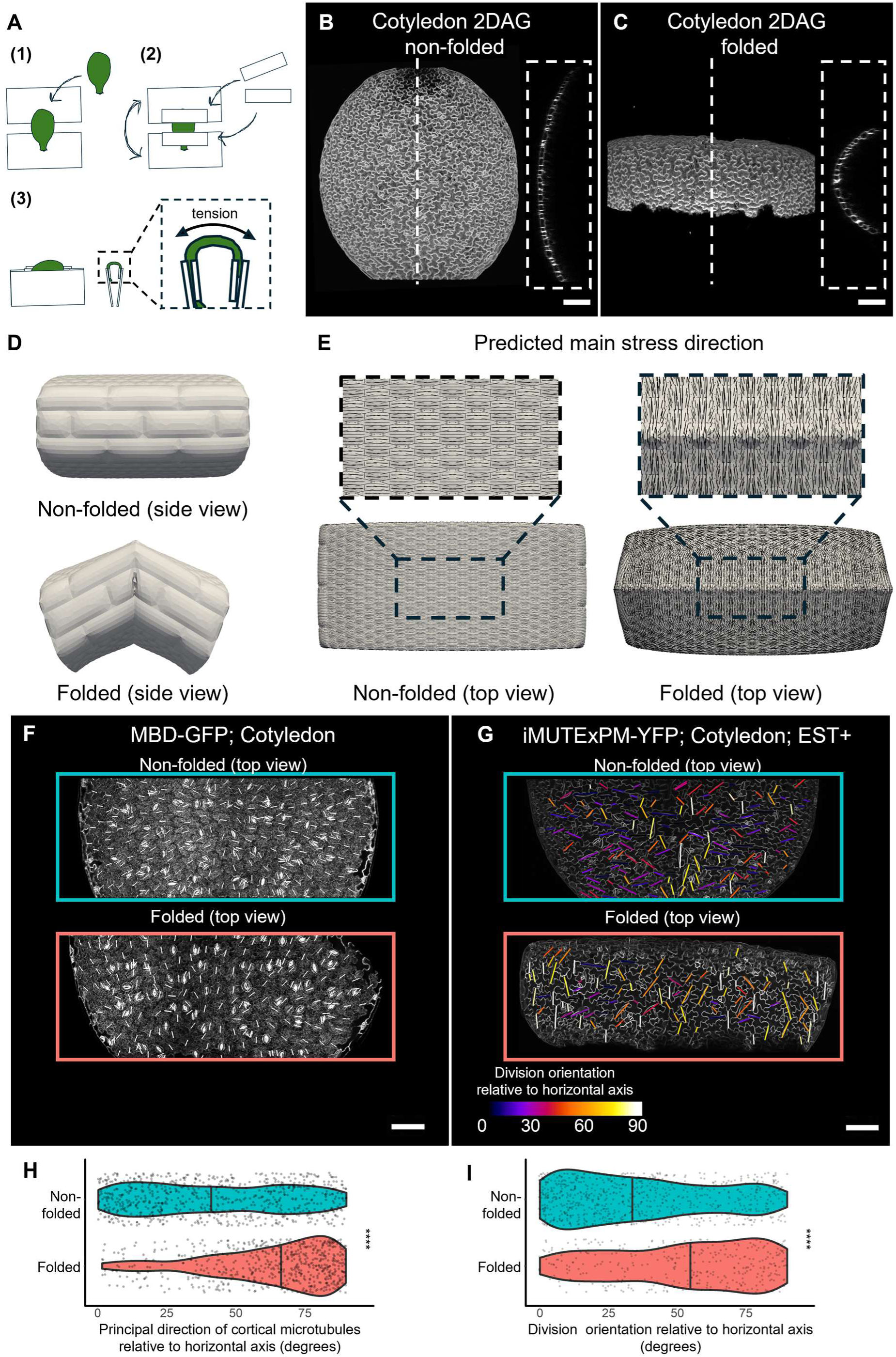
Changes in the tensile stress pattern shift the orientation of stomata divisions. (A) Experimental setup to apply tension on cotyledons: the adaxial side of a cotyledon is stuck on two large pieces of tape (tough tag©) (1), additional pieces of tape are positioned to secure the cotyledon (2), the two large pieces of tape are then folded on top of each other (3) resulting in a folded cotyledon with a predicted tension following the axis of maximal curvature. (B) Non-folded cotyledon 2 DAG, inset shows a transverse optical section along the dashed line (scale bar: 100μm). (C) Folded cotyledon 2 DAG, inset shows a transverse optical section along the dashed line (scale bar: 100μm). (D) Side view of a realistic model of a cotyledon non-folded and folded. (E) Predicted main stress direction on the top side (adaxial) of the non-folded and folded cotyledon model. (F) Organisation of cortical microtubules array (CMT) on non-folded and folded adaxial side of cotyledons (lines show the principal direction of CMT, scale bar:100μm). (G) Orientation of stomata division on non-folded and folded adaxial side of cotyledons (scale bar:100μm). (H) Distributions of CMT orientation relative to a horizontal axis for non-folded and folded cotyledons (data from 2 non-folded and 2 folded cotyledons for a total of 750 and 605 cells analysed). (I) Distributions of stomata division orientation relative to a horizontal axis for non-folded and folded cotyledons.

## Discussion

We have shown that stomata have organ scale alignment in some developmental contexts (Figure 1). Such alignments of features are frequently observed in animal systems, with hairs, bristles, or scales often being aligned with body’s major axis [16]. In animals, these alignments are explained within the context of planar cell polarity, but in plants, although some polarly localized transmembrane proteins have been reported to form organ scale polarity fields, the mechanism at their origins remain elusive [17–19] and there is no evidences that they support the formation of organ scale feature alignments. To try to track down the causative factor aligning stomata division we investigated the role of cell geometry, growth, and mechanical stress (Figure 2 and 4), as they have repeatedly been proposed to play instructive roles in the definition of cell division orientation [5, 7, 8, 20, 21]. Our analyses revealed that stomata divisions were not directed by cell geometry or local growth but instead were more consistently aligned with the predicted pattern of organ scale tensile stress. This result is consistent with previous observations in meristems boundaries domain and stems where divisions are aligned by mechanical stress [20]. While in boundaries and stems the tensile stress pattern aligning division arises from the organ geometry, in the context of the early cotyledon we predict it to arise from the differential growth between the two sides of the cotyledon (Figure 3). A consequence of this differential growth is that more stress relaxation is expected on the fast-growing adaxial side than in the slow-growing abaxial side, which could explain the more disorganised pattern of stomata orientation on the adaxial side (Figure1, and 2) as well as the lower number of cracks observed in the *quasimodo2-2* adaxial cotyledon (Figure 3). It remains to be determined at what stage of development mechanical stress caused by the local cellular geometry may become dominant over the tissue scale stress.

We conclude that the last differentiating division of the stomata lineage can be oriented by tensile stress, this raises the question of how the cell division machinery is informed of the direction of tensile stress. One possibility is that the microtubules (MTs) directly respond and orient divisions according to mechanical stress, the responsiveness of MTs and cell division have been investigated separately [12, 20] but the involvement of MTs to input tensile stress orientation into cell division orientation has not been directly established. Disorganisation of cell division patterns by the application of MTs depolymerising drug Oryzaline in tomato leaves suggests that MTs are involved in this process [22]. Another possibility, which is part of the emerging view that polarity is connecting mechanical stress patterns to morphogenesis [23] is that tension could influence the localisation of transmembrane proteins which will subsequently impact the orientation of cell division. Although it remains under debate, mechanical stress reorientation of polarly localised proteins involved in the stomata lineage has been reported[2, 18]. These proteins (BASL, and BRXL2) have been shown to direct the orientation of cell division by creating a MTs depletion zone in the plasma membrane [24]. In Guard mother cells, the protein OCTOPUS LIKE 2 (OPL2) has been reported to accumulate at both ends of the cell, marking the emplacement of the future division plane. OPL2 is associated with enrichment in MTs and directly interacts with MTs-associated proteins [25]. It is not known if OPL2 responds to mechanical stress, if that were the case, this would make OPL2 an ideal candidate to connect tensile stress patterns with stomata division orientation.

To date, in plants, mechanical stress patterns can only be predicted with models or inferred from MTs orientations[12], tissue deformation upon cut [13] or cracks orientations in mutant lines with adhesion defects [11, 26]. If the alignment of stomata with tensile stress can be generalised in other organs or species, measuring their orientation would provide a recorded history of the tensile stress experimented during the development, in addition, this would make iMUTE inducible line a powerful tool to visualise the tensile stress pattern in various plant organs.

## Material and Methods

### Plant material and growth conditions

All lines are in Columbia ecotype (Col-0), iMUTE line is described in [4], pPDF1::mCitrine-Ka1 (PM-YFP) is from [9], p35S::MBD-GFP (MBD-GFP) has been described in [12], and *quasimodo2-2* in [27]. PM-YFP has been introduced in the iMUTE line by crossing.

Seeds were surface sterilized (10 min in etOH 70%,SDS 0.5%, then washed in etOH 95%, and dried on filter paper) and sown on half-strength MS media (MS/2, 0.8% agar, pH 5.7). Plates were stored at 4 degrees in the dark for 2 to 4 days in order to homogenize germination, then transferred in a grow chamber (20 °C with 16H light/8H dark cycles) in vertical position.

### Sample preparation

Cotyledons were carefully dissected from embryo/seedling at 0, 1 or 2 DAG using fine tweezers under a stereomicroscope, they were then put onto the media surface (MS/2, 0.8% agar,pH5.7, in a Ø35mm petri dish (Grenier bio-one)), and glued to the surface using melted droplets of the same media. For 0DAG samples (germination occurred 24h after stratification in our conditions), mature embryos were extracted from the seed coat by gently squashing the seeds in a droplet of water between two microscopy slides, cotyledons were then dissected and treated as the other samples.

Estradiol inductions were performed using 10 μM β-Estradiol (Sigma Aldrich) containing media.

For timelapse imaging, plates with samples were transferred back to the grow chamber between 2 timepoints.

For dual-view timelapse imaging, samples were prepared as for timelapse imaging, and the samples were carefully flipped upside down using a fine tweezer (Dumont n°5) after the first time point, then re-glued to the surface of the media with a droplet of melted media. After the second timepoint, the plates were transferred back to the grow chamber, and imaged on the following day, after the 3rd timepoint the samples were once again flipped upside down, re-glued to the media, and imaged one last time.

### Mechanical perturbations

For cutting experiments, after dissection, cotyledons suspended in a droplet of water were cut in half with a surgical syringe needle under a stereomicroscope, the cut cotyledons were then placed and glued on the surface of MS/2 media as described above. For needle ablation experiments, cotyledons were prepared as described in the sample preparation section, they were then gently stabbed under a stereomicroscope using a surgical syringe needle.

### Confocal imaging

Petri dishes containing samples were submerged in water for 5 minutes prior to imaging in order to fully hydrate the media and prevent gel swelling induced movements during imaging. All the imaging has been done using long-distance water dipping objectives (HCX APO L 203/0.5-W objective for plasma membrane marker, and HC FLUOTAR L 25x/0,95 W VISIR objective for MBD-GFP) on either an upright Leica SP8 or a Leica Stellaris 5. mCitrine, GFP and Propidium Iodide were excited at 488nm and fluorescence collected between 520nm and 570nm for mCitrine, 498nm and 550nm for GFP, and 575nm and 625nm for Propidium Iodide. All stack of images were acquired with a voxel size no bigger than 0.5x0.5x0.5 μm, for microtubules imaging, the x/y voxel size was reduced to 0.15x0.15 μm.

### Stack focusing imaging

Cotyledons were dissected and positioned on the surface of MS/2 media in petri dishes, they were then imaged using a Keyence VHX-5000 microscope with automated tile scan and extended depth of field function. All samples were imaged using 1000x magnification (VH-Z100R/Z100T wide range zoom lens) with coaxial lighting at low intensity (less than 20%) to prevent sample dehydration.

### Image Analysis

Orientation of stomata relative to proximo distal axis of cotyledons (Figure 1) and division orientation relative to various axes (horizontal axis, cut, or folding axis, for the mechanical perturbation experiment in FigureS2 and Figure 4) has been extracted using homemade macros in ImageJ (https://imagej.nih.gov/ij/). The macro uses annotated images as inputs and outputs heatmaps and .csv files with stomata/cell division orientation. On the raw images, Stomata/new divisions and PDaxis/horizontal axis/cut axis are manually annotated with lines, the macro then allows to segment these lines, compute the relevant angles (e.g stomata orientation v.s PD axis), save data as .csv, and print heatmaps as .tiff.

### MorphoGraphX pipelines

Segmentations, lineage tracking, and growth analyses in MorphographX (https://morphographx.org/; [10, 28]) have been described previously, they are briefly summarized here, for more detailed pipelines refer to [29].

2.5D surface creation, cell segmentation, and lineage tracking: Original stack is blurred (gaussian blur: 0.3/0.3/0.3 μm (x/y/z)), then a solid shape is extracted (edge detect), and a mesh is created (marching cube surface: 5μm), subdivided and smoothed. Signal from the epidermis is projected onto the mesh (project signal 2 to 6 μm) and blurred, cells are manually seeded, and segmentation is done by watershed. The mesh is then refined near cell borders a couple of times.

For lineage tracing, segmented meshes of successive timepoints are loaded in MGX. The mesh of the second timepoint is aligned over the one of the first timepoint, and parent labels are manually transferred using the transfer parent label function.

Computations of axes, and divergence angles measurements:

All of these analyses have been done on the surface of the latest timepoint of timelapse experiments (tn+1) with parent labels from the previous timepoint (tn). A heatmap of cell proliferation was generated to later filter the data to keep only cells that divided one time (clones of 2 cells). Growth axes, clone major axis, and division orientation are extracted using PDG, shape analysis, and fibrils orientation processes. Once these axes have been extracted, divergence angles are computed using the “compute angles” process.

### Light sheet imaging

Lightsheet microscopy was performed using a custom-built laser-scanning light sheet microscope. The design is based on an openspim geometry [30] with dual-side illumination and dual-side detection [31]. Water immersion objectives are mounted horizontally (Nikon 10x, 0.3 NA for excitation, Olympus 20x 1.0 NA for detection) so that their focal points coincide. The sample is attached to a 3d printed holder and suspended from the top between the objectives. For sample placement as well as for imaging, the sample can be moved between the objectives as well as rotated with piezo-driven stage(Nanos LPS-30, Nanos RPS-LW20 for rotation). Image stacks are acquired by moving the sample through the stationary imaging plane. A 488nm laser (Omicron LuxX 488-200) was used for excitation. The fibre output was collimated, galvo scanned (Galvo system: Thorlabs GVSM002-EC/M) and magnified resulting in a scanned light sheet with typical FWHM < 5μm. Only one of the two possible sCMOS cameras (Hamamatsu Orca Flash 4) with 6.5x6.5 μm² pixel size was used for detection. A bandpass filter (Chroma ET525/50m) allows the recording of the specific fluorescence band. The microscope is controlled by a custom software developed in LabVIEW (National Instruments). Data was streamed to disk and converted to TIFF files directly after acquisition resulting in image voxel sizes of 1 μm³.

The plants were attached to the holder, lowered in between the four objectives. They were illuminated from two sides to minimise shadowing artifacts. Typical excitation powers set in software were 3% for 488nm excitation. The camera exposure time was set to 100ms per plane. Multiple positions were recorded in a grid pattern and stitched in Fiji [32] using the grid/collection stitching plugin [33] before further analysis.

### Modelling

A realistic cotyledon mesh was generated using a custom code (https://gitlab.developers.cam.ac.uk/slcu/teamsr/meshing_code) and a Delaunay-based unstructured mesh generation algorithm from Darren Engwirda [34]. The cotyledon mesh contains an adaxial, abaxial epidermis mesophyll and palisade. All widths and heights of the cells were based on actual measurements from a cotyledon 1DAG (Figure 3F). We also kept the curvature of the cotyledon and the aspect ratio of the cotyledon the same to get an accurate geometry. To remove the influence of cell shape we used square cells which have isotropic stress when isolated.

All meshes were created so that each connection between two nodes were approximately 0.3 *μm long*. This mesh was then converted into a compatible form to inflate in *Tissue*, software developed by the Jӧnsson group and further developed for 3D by the Robinson group [12, 35–37].

*Tissue* uses a finite element method approach to approximate solutions to the hyperelastic continuous mechanical equations over the discretized mesh surface, which are being stretched by a normal (outward from each cell) pressure force applied to each mesh triangle. It uses the ‘Triangular Bi-Quadratic Springs’ (TRSB) method, which uses a St Venant Kirchoff formulation and approximates it using biquadratic springs, which resist triangle edge and inner angle deformation [38] and allows us to simulate cell walls efficiently. In more detail, the St Venant Kirchoff formulation stress-strain relation for an isotropic material is,

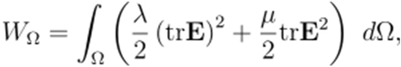

where *W* is total strain energy in a domain *Ω*, **E** the Green-Lagrange strain tensor and *λ* and *μ* are the Lame coefficients and defined in plane elasticity as 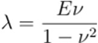 and 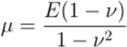 with *E* being the Young’s modulus and 𝜈 the Poisson coefficient. Using TRBS, *W* is approximated over a triangle, *T,* as

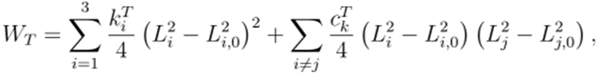

where *k_i_^T^* and *c_i_^T^* are the tensile and angular stiffness of the biquadratic springs and *L_i_* and *L_i,0_* are the current and resting lengths of edge *i* in the triangle. These stiffnesses are defined as

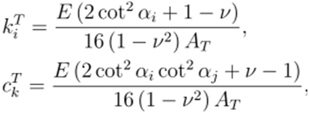

where *α*_i_ is the angle opposite edge *i* in the resting triangle configuration and *A_T_* the resting triangle area. A force on each node can then be calculated from this strain energy as a result of the stretching triangles. Additionally, deriving the stress tensor for each triangle allows us to find the maximum stress magnitude and direction from the maximum corresponding eigenvector and eigenvalue.

The pressure force is applied to all the triangles proportional to their area in the outward normal direction. A solution to the simulation is found when the system has reached mechanical equilibrium, i.e. when the pressure forces on each node acting on them from their surrounding triangles are balanced with their surrounding stretched and strained triangles, such that the total force is now *0*. We find this mechanical equilibrium using the Newton-Raphson method [39] and setting the tolerance to *10^-3^*.

The growth simulations were performed by first inflating to mechanical equilibrium and saving their state. The system was then allowed to grow using a spring with a resting length that is permitted to extend, a growth formulation used previously to model plants [40, 41]. Specifically, the resting length, *L^t+δt^_i,0_* of the element *i* at time *t+ δt* (with time step size *δt*) is updated using it resting length in the previous time step, *L^t^_i,0_* plus its defined extension, which is proportional to their strain and their current length. This formulation takes the form,

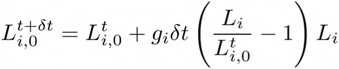

where *g_i_* is the extensibility rate and *L_i_* the current length of that edge [12, 42].

All simulations were inflated with a turgor pressure of 0.1 MPa, Young’s modulus of 100 MPa and a Poisson’s ratio of 0.2. The values chosen for the turgor pressure and the walls Young’s modulus are in line with previous studies [15, 43]. The anticlinal cell walls parallel to the cotyledon long axis were set to 500MPa to help preserve the aspect ratio of the cotyledon as it grew. The growth rates were set between 1 at the top of the tissue and 0 at the bottom. This was set to replicate the difference in magnitude of the growth between the abaxial and adaxial sides as seen in the real data (Figure 3D and E). The model makes the simplifying assumption that the growth rate decreases linearly through the inner layers, is isotropic, and was scaled to be between 0 and 1. For the three points bending we applied forces along the horizontal length of our mesh. These forces had direction and magnitude for each node of [0 -197.28 - 32.88], [0 -197.28 +32.88] and [0 257.4134759 0] for the front, back and top respectively set so that the forces balance out (where the y direction is from top to bottom, x the horizontal axis and z the PD axis).

The black lines on the Figures are the principal stress direction from the Cauchy stress tensor.

## Authors contributions

L.S and S.R designed the project, L.S performed the experiments and analysed the data with the help of L.B, E.T.S developed and ran the simulations, M.O.L built the light-sheet microscope and performed the light-sheet imaging, L.S and S.R wrote the manuscript with inputs from E.T.S and M.O.L, S.R supervised the project and acquired funding.

## Acknowledgments

S.R. was supported by University Research Fellowships 2018 (URF\R1\180196). S.R. and by the Gatsby Charitable Foundation (G101113). L.S was supported by The Leverhulme Trust (RPG-2022-111). We acknowledge the support of the professional services at SLCU for their help with this project.

## Supplemental figures

**Figure S1:**
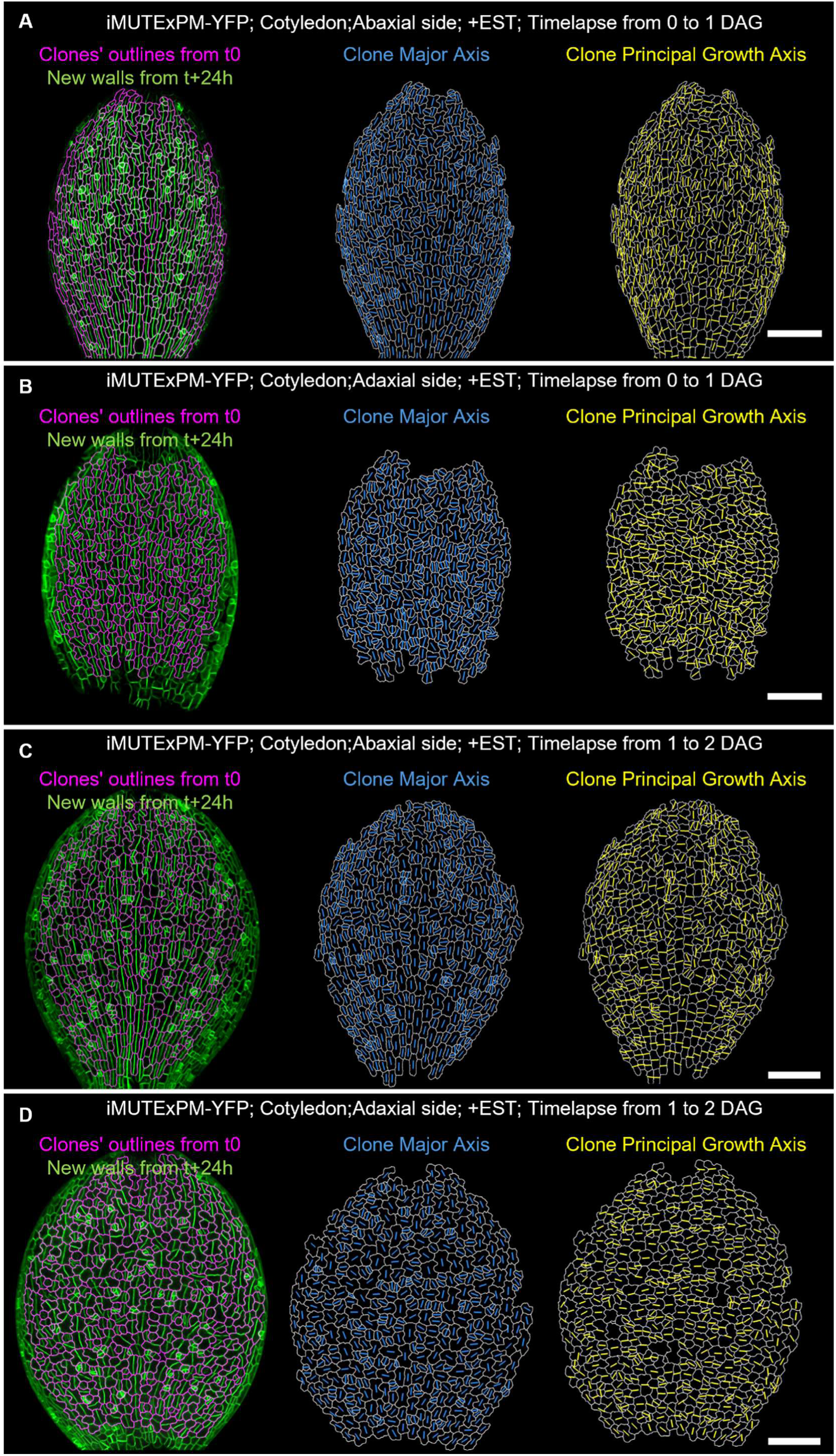
Clone principal growth axis and clone major axis for iMUTExPM-YFP timelapses. (A) Stomata differentiating divisions induced between 0 and 1 DAG on the abaxial side of iMUTExPM-YFP cotyledons (Clones from t0 are outlined in magenta, the surface at t+24h highlighting the new walls is in green), Clone major axis in blue and clones principal growth axis in yellow (scale bar:100μm). (B) Stomata differentiating divisions induced between 0 and 1 DAG on the adaxial side of iMUTExPM-YFP cotyledons (Clones from t0 are outlined in magenta, the surface at t+24h highlighting the new walls is in green), Clone major axis in blue and clones principal growth axis in yellow (scale bar:100μm). (C) Stomata differentiating divisions induced between 1 and 2 DAG on the abaxial side of iMUTExPM-YFP cotyledons (Clones from t0 are outlined in magenta, the surface at t+24h highlighting the new walls is in green), Clone major axis in blue and clones principal growth axis in yellow (scale bar:100μm). (D) Stomata differentiating divisions induced between 1 and 2 DAG on the adaxial side of iMUTExPM-YFP cotyledons (Clones from t0 are outlined in magenta, the surface at t+24h highlighting the new walls is in green), Clone major axis in blue and clones principal growth axis in yellow (scale bar:100μm).

**Figure S2:**
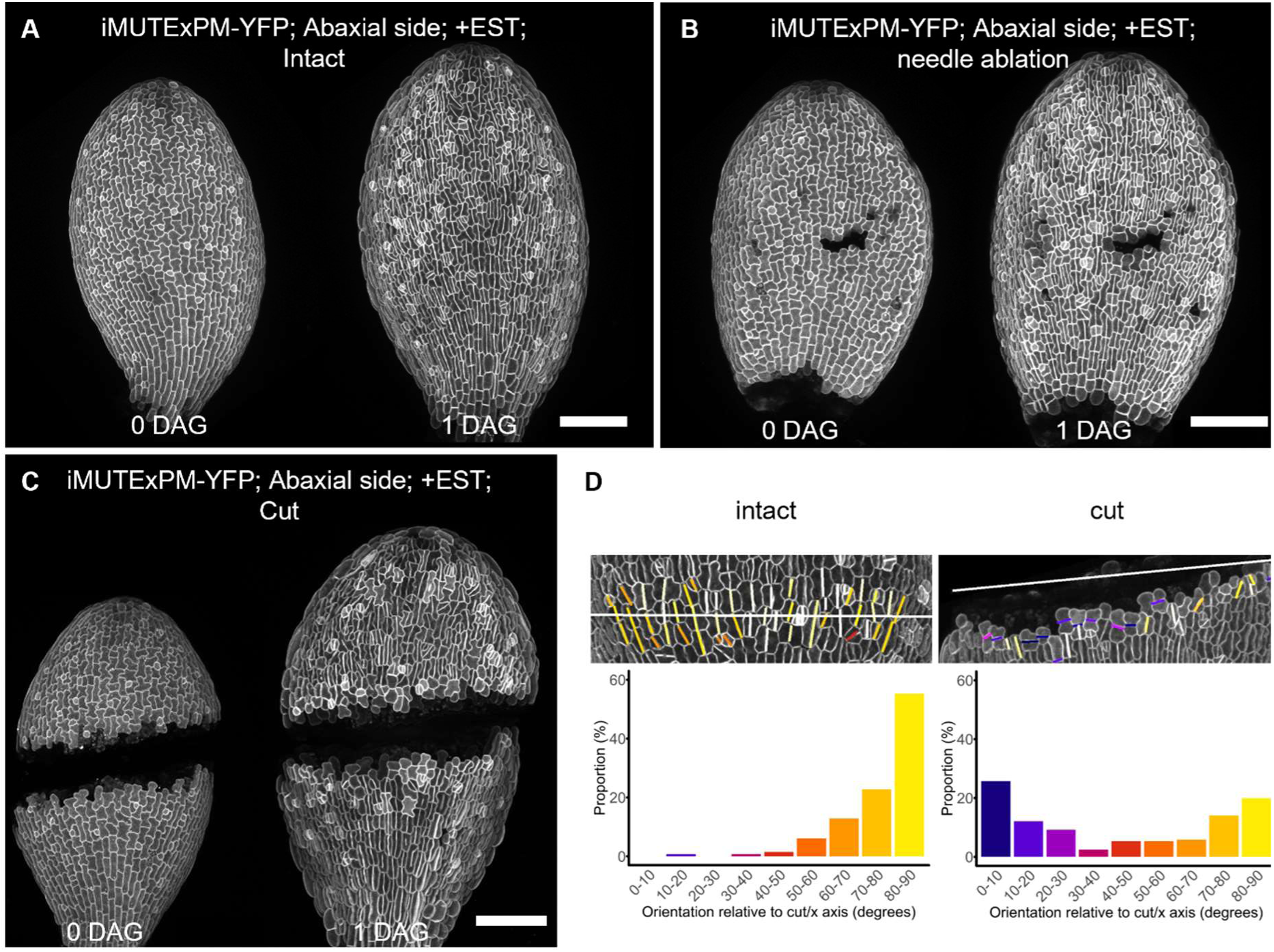
Mechanical perturbations changes the orientation of stomata differentiating divisions. (A) Snapshots of abaxial side of intact iMUTExPM-YFP cotyledons at 0 and 1 DAG with division induced between the two timepoints (scale bar 100μm). (B) Snapshots of abaxial side of iMUTExPM-YFP cotyledons at 0 and 1 DAG after needle ablation at 0DAG, and division induced between the two timepoints (scale bar 100μm). (C) Snapshots of abaxial side of iMUTExPM-YFP cotyledons at 0 and 1 DAG after needle cutting at 0DAG, and division induced between the two timepoints (scale bar 100μm). (D) Quantification of division orientation relative to either a horizontal axis of the axis of the cut for the cotyledon needle cutting experiment in (C).

